# Effects of abiotic environment on invertebrate herbivory depend on plant community context in a montane grassland

**DOI:** 10.1101/2022.04.19.488732

**Authors:** Fletcher W. Halliday, Seraina L Cappelli, Anna-Liisa Laine

**Affiliations:** Department of Evolutionary Biology and Environmental Studies, University of Zurich, 8057, Zurich, CH; Research Centre for Ecological Change, Organismal & Evolutionary Biology Research Programme, Faculty of Biological and Environmental Sciences, PO Box 65, FI-00014 University of Helsinki, Finland

**Keywords:** biodiversity, herbivory, community structure, elevation, climate change

## Abstract

Invertebrate herbivores are important and diverse, and their abundance and impacts are expected to undergo unprecedented shifts under climate change. Yet, past studies of invertebrate herbivory have documented a wide variety of responses to changing temperature, making it challenging to predict the direction and magnitude of these shifts. One explanation for these idiosyncratic responses is that changing environmental conditions may drive concurrent changes in plant communities and herbivore traits. Thus, the impacts of changing temperature on herbivory might depend on how temperature combines and interacts with characteristics of plant communities and the herbivores that occupy them. Here, we test this hypothesis by surveying invertebrate herbivory in 220, 0.5 meter-diameter herbaceous plant communities along a 1101-meter elevational gradient. Our results suggest that increasing temperature can drive community-level herbivory via at least three overlapping mechanisms: increasing temperature directly reduced herbivory, indirectly affected herbivory by reducing phylogenetic diversity of the plant community, and indirectly affected herbivory by altering the effects of functional and phylogenetic diversity on herbivory. Consequently, increasing functional diversity of plant communities had a negative effect on herbivory, but only in colder environments while a positive effect of increasing phylogenetic diversity was observed in warmer environments. Moreover, accounting for differences among herbivore feeding guilds considerably improved model fit, because different herbivore feeding guilds varied in their response to temperature and plant community composition. Together, these results highlight the importance of considering both plant and herbivore community context in order to predict how climate change will alter invertebrate herbivory.

## Introduction

Outbreaks of invertebrate herbivore pests are becoming increasingly common, with much of this increase attributed to global climate change (Deutsch et al. 2018, Science). Yet, attributing the extent of invertebrate herbivory to climate change is complicated by the fact that increasing environmental temperature can affect herbivory through a variety of mechanisms (Moreira *et al*. 2018). Increasing temperature can directly impact herbivore growth, survival and reproduction (Huey & Kingsolver 1989), but can also indirectly influence herbivory via two general pathways. First, changing temperature can alter the composition and diversity of plant communities that herbivores rely on, thereby determining which plant-herbivore interactions occur in a given place or time (**mediation**) (Hülber *et al*. 2015; Warner *et al*. 2021). Second, changing temperature can alter the role that plant communities play in driving herbivory (**moderation**) (Field *et al*. 2020), for example by changing the direction or magnitude of particular plant-herbivore interactions (Wolinska & King 2009; Descombes *et al*. 2017; Pellissier *et al*. 2018; Galmán *et al*. 2021). Thus, predicting how climate change will affect invertebrate herbivory will require an understanding of how multiple factors combine and interact with one another across environmental gradients (Fig. 1).

**Figure 1.**
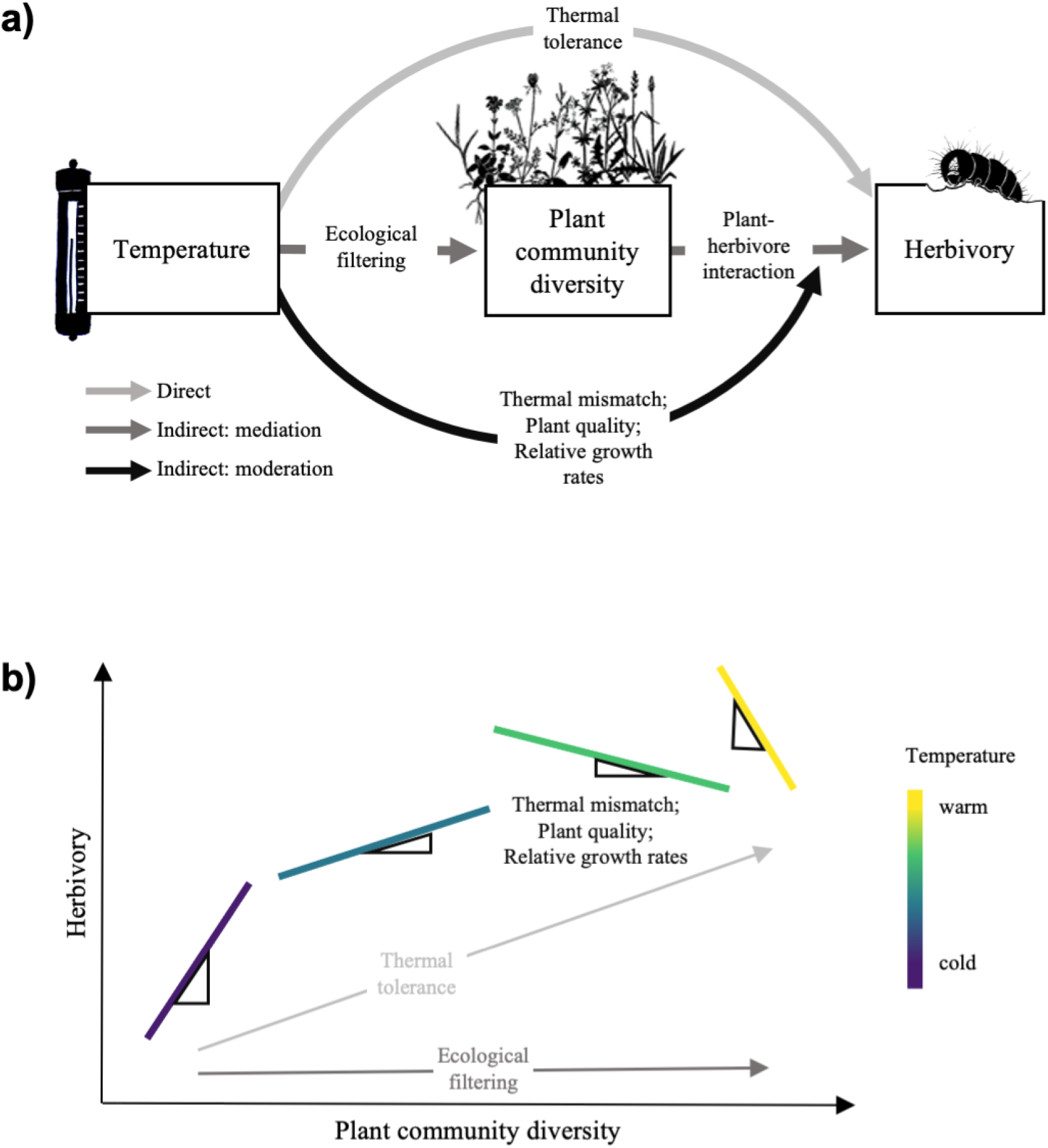
Hypothesized relationships among temperature, plant community diversity, and invertebrate herbivory. A) Increasing environmental temperatures can directly influence herbivory (e.g., by acting on thermal tolerance of invertebrates, illustrated in light grey), and indirectly influence herbivory, both by altering characteristics of plant community diversity (e.g., via ecological filtering, mediation, illustrated in dark grey), and by modifying plant-herbivore interactions (e.g., via thermal mismatch of plants and herbivores, moderation, illustrated in black). B) A hypothetical example of a second-stage moderated mediation, in which temperature influences herbivory via all three pathways simultaneously.

The direct impacts of environmental temperature on the performance of invertebrate herbivores are well studied. Traditionally, cooler climates are expected to exhibit lower overall levels of invertebrate herbivory (Bale *et al*. 2002; Metcalfe *et al*. 2014; Kozlov *et al*. 2015; but see Moreira *et al*. 2015; Galmán *et al*. 2018). Cooler temperatures can directly limit the performance of invertebrate herbivores by reducing herbivore metabolic activity, growth rate, and reproductive output (e.g., thermal tolerance; Huey & Kingsolver 1989; Kingsolver *et al*. 2013; Andrade *et al*. 2020). Cooler temperatures can also affect herbivores by reducing plant defense compounds, palatability, and tissue nutrient concentrations (Andrade *et al*. 2020; Hamann *et al*. 2021). Thus, changing environmental conditions can have dramatic impacts on the performance of invertebrate herbivores (Hamann *et al*. 2021; Kharouba & Yang 2021) (Fig. 1 light grey arrows).

In complex ecological communities, herbivory often depends on the distributions and traits of plant species (Haddad *et al*. 2001; Scherber *et al*. 2006; Barbosa *et al*. 2009; Loranger *et al*. 2014; Underwood *et al*. 2014; Ebeling *et al*. 2018; Gardarin *et al*. 2018; Field *et al*. 2020), which themselves are sensitive to environmental conditions (Whittaker 1956; Díaz & Cabido 1997; Lavorel & Garnier 2002; McMahon *et al*. 2011; Sundqvist *et al*. 2013; Reich 2014; Harrison *et al*. 2020), generating an indirect pathway through which environmental temperature can affect herbivory (mediation, Fig. 1 dark grey arrows) (Pitteloud *et al*. 2020; Warner *et al*. 2021). Increasing temperature is often associated with changes in plant taxonomic, functional, and phylogenetic diversity (e.g., via ecological filtering; Chapin *et al*. 1994; Klein *et al*. 2004; Gedan & Bertness 2009; Coyle *et al*. 2014; Zhu *et al*. 2020), though the direction and magnitude of effects across dimensions of biodiversity and ecological systems is far from certain (Shi *et al*. 2015; Chase *et al*. 2019; Dornelas *et al*. 2019). These multiple dimensions of plant diversity can, in turn, influence plant-herbivore interactions through associational effects (Barbosa *et al*. 2009; Underwood *et al*. 2014; Field *et al*. 2020), which can determine the ability of a plant community to provide resources and habitat to different arthropod groups (Gardarin *et al*. 2018). In two pioneering biodiversity experiments, increasing taxonomic diversity consistently increased the abundance of invertebrate herbivores (Haddad *et al*. 2001; Scherber *et al*. 2006; Loranger *et al*. 2014; Ebeling *et al*. 2018), while increasing functional diversity reduced herbivore abundance in one study (Haddad *et al*. 2001), but increased herbivore abundance in another (Scherber *et al*. 2006). Increasing phylogenetic diversity could increase herbivory (e.g., Parker *et al*. 2012; Egorov *et al*. 2017) by providing a greater diversity of resources to herbivores, particularly when plant palatability is phylogenetically conserved. However, the opposite pattern could emerge if increasing phylogenetic diversity “dilutes” the availability of resources for specialists as it can for the transmission of plant parasites (e.g., Halliday *et al*. 2019).

The effects of plant communities on community-level herbivory might also be sensitive to local abiotic conditions (Richardson *et al*. 2002; Rasmann *et al*. 2014; Hülber *et al*. 2015), thereby generating a second indirect pathway through which environmental temperature can affect herbivory (moderation, Fig. 1 black arrow). Specifically, the effects of temperature on plant quality (e.g., plant defense, palatability, tissue nutrients), growth rates, or phenology can vary considerably among taxa from different clades or functional groups (Herz *et al*. 2017; Roybal & Butterfield 2019; Kuppler *et al*. 2020; Hamann *et al*. 2021; Sanczuk *et al*. 2021; Sandel *et al*. 2021) (but see Midolo *et al*. 2019). Variation in how taxa within a community respond to changing temperature could result in thermal mismatch between the plant community and insect herbivores, and alter the role of phylogenetic or functional diversity in driving herbivory across temperature regimes. Thus, characteristics of plant communities associated with higher levels of invertebrate herbivory in one environment might not predict invertebrate herbivory under environmental change.

Finally, how these diverse and interacting drivers of herbivory play out in nature might also depend on characteristics of the herbivores themselves (Tscharntke & Greiler 1995; Bale *et al*. 2002). Specifically, differences among invertebrate herbivore feeding guilds (e.g., invertebrates that are free living and feed externally on plant leaves vs. those that live and feed inside of plant leaves) can determine the impact of changing environmental conditions on herbivory (Anstett *et al*. 2014; Andrade *et al*. 2020). Yet, the degree to which herbivore feeding guilds influence responses to interacting biotic and abiotic factors remains unknown. If certain feeding guilds that respond more strongly to plant taxonomic, phylogenetic, or functional diversity differ in their sensitivity to the abiotic environment, then this could drive complicated interactions in the relationship between biotic and abiotic drivers of herbivory across changing environmental conditions.

In this study, we apply a simple ecological framework (Fig. 1) to understand the response of community-level herbivory to changing temperature and plant diversity along a 1101 m elevation gradient in Southeastern Switzerland. Our results suggest that increasing temperature can drive community-level herbivory via three non-exclusive mechanisms: temperature modifies herbivory directly, but also by changing multiple dimensions of plant community diversity, and by altering the role that plant communities play in driving herbivory. However, these results vary among herbivore feeding guilds, and incorporating herbivore feeding guild into our analysis improves the explanatory power of our framework considerably. Together, these results highlight the context-dependence that is inherent in understanding how changing temperature affects invertebrate herbivory. Nevertheless, because this framework is grounded in fundamental ecological principles, our results also indicate that implementing a simple ecological framework in the context of herbivore feeding guilds and the plant communities that they occupy may be sufficient for predicting how herbivory and herbivore communities will respond to increasing environmental temperatures associated with future climate change.

## Results

To evaluate how plant communities influence invertebrate herbivory across environmental gradients, we surveyed 220, 0.5 m-diameter vegetation communities (i.e., small plots), that were established in four meadows along a 1101 m elevational gradient as part of the Calanda Biodiversity Observatory (CBO; Halliday *et al*. 2021). We then recorded the identity and visually quantified the percent cover of all plant taxa in each vegetation community. Vegetation communities in the CBO consist mostly of perennial herbs and grasses that tolerate grazing. In the CBO, mean soil, soil surface, and air temperature all strongly and consistently decreased with increasing elevation, while mean soil moisture was uncorrelated with elevation (Halliday *et al*. 2021). Here, we report analyses using air temperature, and because soil moisture was uncorrelated with elevation, host community structure, and disease (Halliday *et al*. 2021), and was similarly unrelated to invertebrate herbivory, this factor was omitted from analyses.

### Abiotic drivers of plant taxonomic, functional, and phylogenetic diversity

As observed in a previous analysis of the CBO, species richness in the small plots was highly variable (7–30 species) and increased with elevation, while temperature declined (Halliday *et al*. 2021) (Table S1; Figure S1a).

Community functional diversity was calculated independent of species richness by computing how the abundance of species in a community were distributed within the volume of functional trait space occupied by those species, calculated with traits extracted from the TRY database (Kattge *et al*. 2020) using a measurement called functional divergence (FDiv; Villéger *et al*. 2008). This measure of functional diversity was estimated using seven foliar functional traits (specific leaf area, carbon-to-nitrogen ratio, leaf chlorophyll content, leaf lifespan, leaf nitrogen, leaf phosphorus, and photosynthetic rate) as well as plant height and seed mass, all of which have been shown to underlie important tradeoffs related to species growth, reproduction, competition, and defense against natural enemies (Westoby *et al*. 1996; Wright *et al*. 2004; Reich 2014; Díaz *et al*. 2016). Community functional diversity was independent of species richness (r = −0.04). Unlike species richness, functional diversity was highest at low elevation and declined as elevation increased and temperature declined, though this effect was only weakly supported by the statistical model and explained little of the overall variation in community functional diversity (p = 0.07, Marginal R^2^ = 0.05; Conditional R^2^ = 0.56; Table S1; Fig S1b).

Community phylogenetic diversity was calculated independent of community functional diversity and species richness by computing the degree to which plant communities were more or less phylogenetically diverse than random, given the number of plant species and their relative abundance. Across all sites, plant communities were commonly phylogenetically clustered, meaning that plant species were more closely related than would be expected based on the number of plant species and their relative abundance (Fig S1). Community phylogenetic diversity was independent of species richness (r = 0.17) and functional diversity (r = 0.007) and decreased (i.e., became more phylogenetically clustered) with increasing air temperature (p = 0.003; Marginal R^2^ = 0.16; Conditional R^2^ = 0.70; Table S1; Fig S1c).

### Biotic and abiotic drivers of invertebrate herbivory

We quantified herbivory by visually surveying leaves for damage, and recording the percent of leaf area that had been consumed by invertebrate herbivores following the plant pathogen and invertebrate herbivory protocol in Halbritter et al (2020). In order to understand how the abiotic environment and plant community structure combine and interact to influence invertebrate herbivory, we constructed a piecewise structural equation model (SEM). This model explores whether changes in invertebrate herbivory along the elevational gradient are (partially) mediated by changes in plant taxonomic, phylogenetic, and functional diversity, as well as whether the effect of taxonomic, phylogenetic, and functional diversity on invertebrate herbivory might change as a function of increasing temperature (i.e., are moderated by temperature). The data were well fit by this model (Fisher’s C = 13.14; P-value = 0.515; 14 degrees of freedom, Table S2; Fig. 2). The model leverages the strong, negative effect of elevation on air temperature (−1.04, p < 0.001, R^2^ = 0.89) to compare separate pathways through which changing temperature can affect herbivory. The model included the previously measured strong declines in host plant richness (−0.32, p = 0.005) and phylogenetic diversity (−0.19, p = 0.009) and weak (and statistically not significant) increase of host functional diversity (0.18, p = 0.09) with increasing temperature (and thus decreasing elevation, Fig 2).

**Figure 2.**
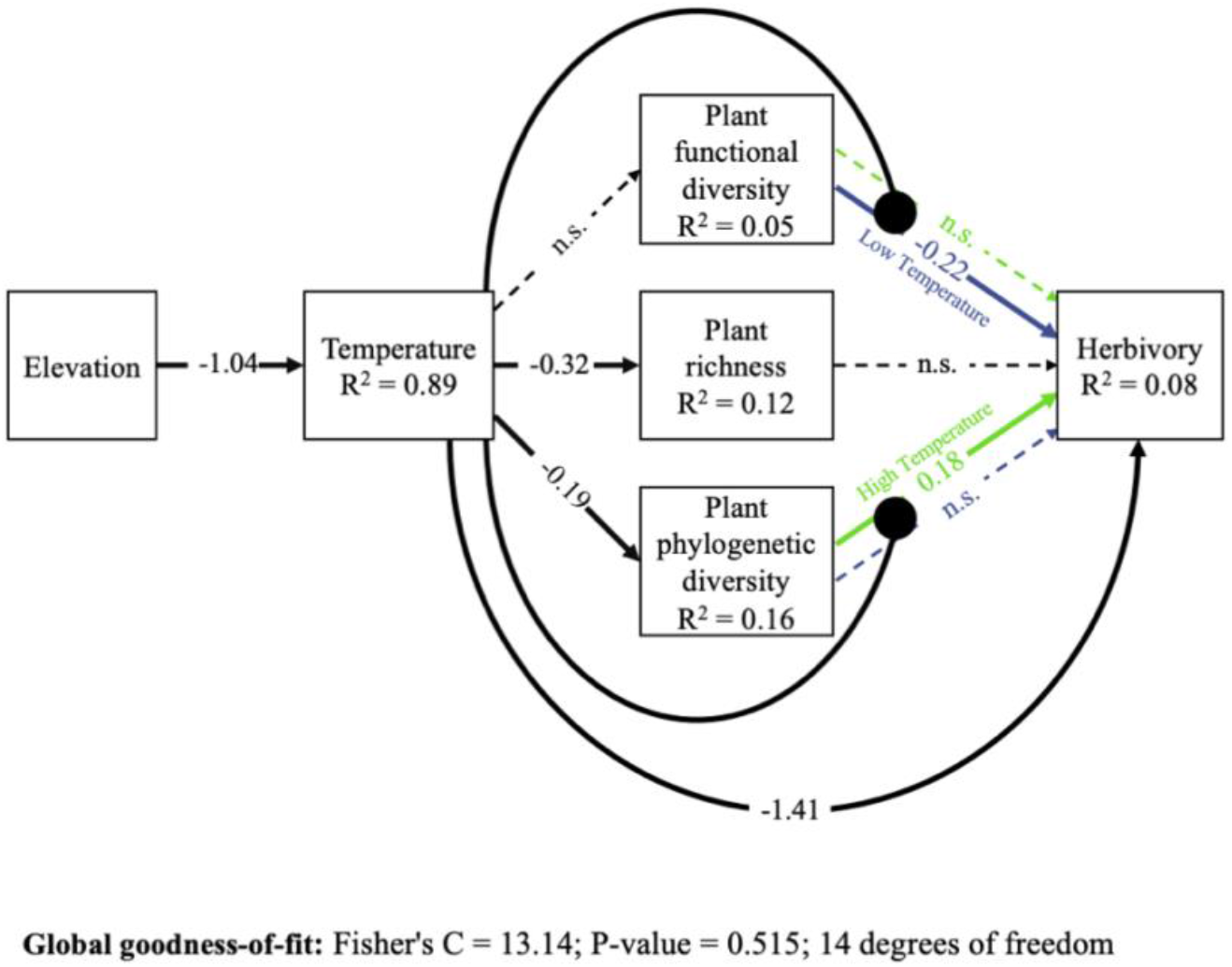
Results from the piecewise structural equation model. All coefficients are scaled by the ratio of the standard deviation of x divided by the standard deviation of y (i.e., standardized estimates). Coefficients that were not supported by the model (p < 0.05) are illustrated using a dashed line. Colors are drawn to highlight the statistical interactions between host community functional and phylogenetic diversity and temperature. The High and Low Temperature coefficients are estimated with the reference temperature set to 1 standard deviation above and below the mean temperature, respectively. All other coefficients are estimated from a model using mean-centered values for temperature. Correlations between errors were not supported by the model and are not shown. A full list of model coefficients is provided in Table S3. Higher air temperature, associated with lower elevation, affected herbivory through four non-mutually exclusive pathways: directly via abiotic constraints, and indirectly via shifting host phylogenetic diversity as well as by altering the relationship between functional and phylogenetic diversity and herbivory.

As expected, temperature drove invertebrate herbivory through a variety of direct and indirect pathways. First, increasing temperature directly reduced herbivory (−1.41, p = 0.03, Fig. 2), contrary to the expectation that invertebrate herbivores would increase their activity and cause greater herbivore damage with increasing temperature. Second, increasing temperature indirectly reduced herbivory via its negative effect on host phylogenetic diversity. Finally, increasing temperature indirectly affected herbivory by altering the effects of both functional diversity and phylogenetic diversity on herbivory.

This change in the relationship between diversity and herbivory with increasing temperature resulted in more positive diversity effects on herbivory with increasing temperature. Specifically, the effect of functional diversity on herbivory increased from negative to undetectable (moderation path p = 0.047, Fig 2 and 3a), which means that communities in low temperature environments experienced lower rates of herbivory (−0.22, p = 0.011 at 1SD below mean temperature) with increasing functional diversity, but that effect weakened (−0.11, p = 0.065 at mean temperature) and ultimately disappeared as temperature increased (0.0008, p = 0.99 at 1SD above mean temperature). The effect of phylogenetic diversity on herbivory changed from undetectable (−0.17, p = 0.19 at 1SD below mean temperature) to positive (0.18, p = 0.026 at 1SD above mean temperature) with increasing temperature (moderation path p = 0.033, Fig 2 and 3b). In other words, communities in low temperature environments had herbivory rates that were independent of phylogenetic diversity, while at high temperatures, herbivory increased with increasing phylogenetic diversity. These combined effects resulted in a change from a negative diversity effect on herbivory based on functional diversity in colder, high-elevation environments to a positive diversity effect on herbivory based on phylogenetic diversity in warmer, low-elevation environments. In contrast to effects of functional and phylogenetic diversity on herbivory, there was only week evidence for a negative relationship between plant richness and herbivory (−0.11, p = 0.065), and no evidence for any statistical interaction between plant richness and temperature (0.42, p =0.13).

**Figure 3.**
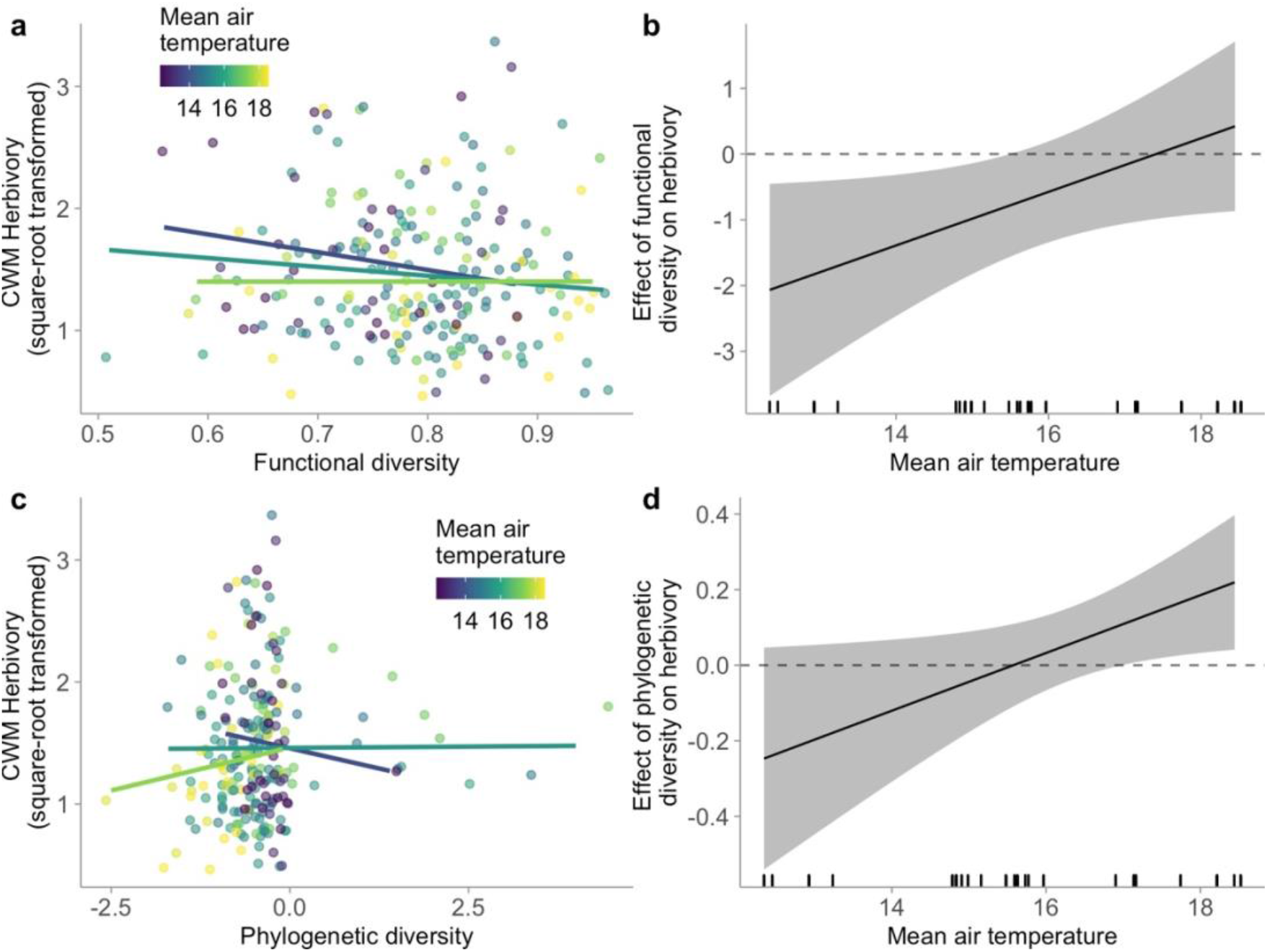
Results of the SEM testing how the response of invertebrate herbivory to community functional diversity (panels a and b) and community phylogenetic diversity (panels c and d) depends on air temperature. In panels a and c, the lines represent the model-estimated effect of functional and phylogenetic diversity, respectively, estimated at one standard deviation above the mean (light green), one standard deviation below the mean (purple), and at the mean temperature (blue). Points represent the raw data colored by air temperature of the site. Increased temperature altered the effect of functional diversity (a) and phylogenetic diversity (b) on insect herbivory (change in slopes) and in addition decreased phylogenetic diversity (change in range along x-axis). In panels b and d, the black line represents the model-estimated mean effect of functional and phylogenetic diversity, respectively, on herbivory (i.e., the slope of the lines in (a) and (c) as a function of increasing air temperature. The grey ribbon is the model-estimated 95% confidence interval.

Overall, instead of observing a net decrease in herbivory with increasing elevation that we expected, this context dependence in the relationships between plant community structure and herbivory generated a net increase in herbivory with increasing elevation, with low overall explanatory power (Marginal R^2^ = 0.08, Conditional R^2^ = 0.31), suggesting a large amount of unexplained variation in herbivory along the elevational gradient.

### Differences in the biotic and abiotic drivers of herbivory among herbivore feeding guilds

We next assessed the degree to which functional, phylogenetic, and taxonomic drivers of herbivory and their interaction with changing temperature might vary among different herbivore feeding guilds using a linear mixed model with a multivariate response (following Halliday *et al*. 2017). We grouped herbivory into four feeding guilds modified from the plant pathogen and invertebrate herbivory protocol in Halbritter et al (2020): chewing, skeletonizing, thrips, and leaf mining (Fig S2). Chewing was identified as missing areas of leaf tissue, mostly caused by orthopterans and lepidopterans. Skeletonizing was identified as the removal of leaf epidermal tissue, mostly caused by mollusks and coleopterans. Damage by thrips was identified as small white patches associated with insects in the order Thysanoptera. Leaf mining was identified as unbroken trails on the leaf surface, caused by a diverse group of invertebrates including lepidopterans, coleopterans, hymenopterans, and dipterans. Galling, which is identified as abnormal growths in the leaf tissue often caused by dipterans and hymenopterans, was observed infrequently and therefore not included in the analysis. Although different invertebrate herbivores can cause highly similar types of damage (Kozlov *et al*. 2016), we expected these four feeding guilds to exhibit different relationships with biotic and abiotic drivers of herbivory (Tscharntke & Greiler 1995; Bale *et al*. 2002; Thaler *et al*. 2012; Anstett *et al*. 2014; Andrade *et al*. 2020), and therefore that taking feeding guild into account might improve the explanatory power of biotic and abiotic drivers of herbivory across the gradient. Consistent with this expectation, the impact of host-community level drivers on herbivory and their interaction with temperature varied among herbivore feeding guilds (Table 1, Fig 4), and incorporating feeding guild into our model of herbivory considerably improved the explanatory power of the model (Marginal R^2^ = 0.385, Conditional R^2^ = 0.388).

**Table 1.** Results of Type II Analysis of Deviance tests on the multivariate response regression quantifying the effects of air temperature, plant community diversity, and their interactions, on community-weighted mean herbivory grouped into four feeding guilds.

| Response | Chisq | Df | P |
| --- | --- | --- | --- |
| Herbivore Feeding Guild | 1651.643 | 4 | <0.0001 |
| Feeding Guild × Air Temperature | 9.545 | 4 | 0.0488 |
| Feeding Guild × Species Richness | 13.430 | 4 | 0.0094 |
| Feeding Guild × Functional Diversity | 11.071 | 4 | 0.0258 |
| Feeding Guild × Phylogenetic Diversity | 22.321 | 4 | 0.0002 |
| Feeding Guild × Temp × Richness | 11.372 | 4 | 0.0227 |
| Feeding Guild × Temp × Functional | 12.094 | 4 | 0.0167 |
| Feeding Guild × Temp × Phylogenetic | 19.036 | 4 | 0.0008 |

**Figure 4.**
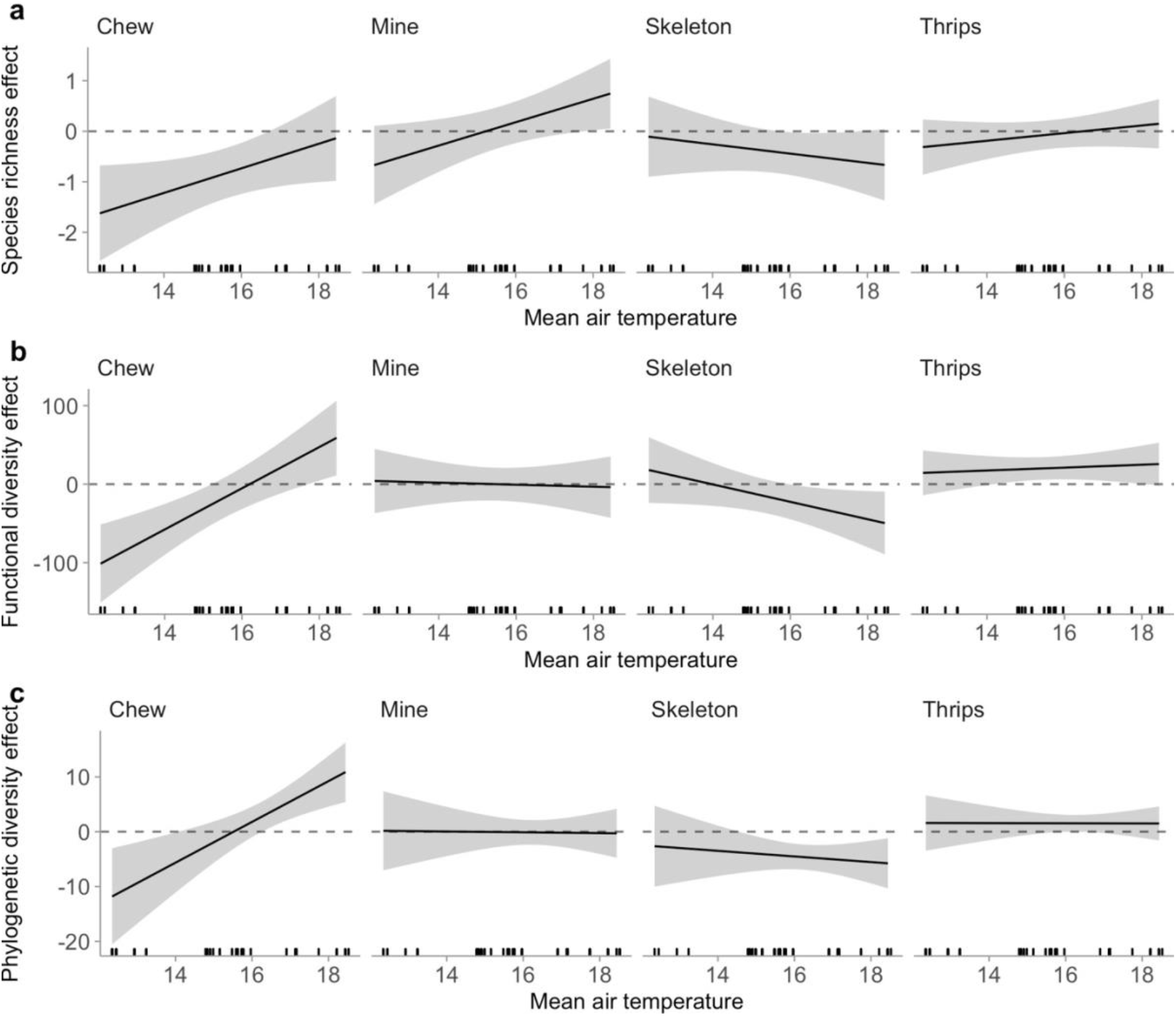
Results from a multivariate response regression, showing that the interaction between taxonomic, functional, and phylogenetic diversity and mean air temperature depends on herbivore feeding guild. The black lines represent the model-estimated mean effect of (a) species richness, (b) functional diversity, and (c) phylogenetic diversity, on herbivory, as a function of increasing air temperature. The grey ribbon is the model-estimated 95% confidence interval.

Chewing and skeletonizing herbivory exhibited distinct responses to biotic and abiotic drivers. Chewing declined with increasing functional and phylogenetic diversity at high elevation sites, characterized by low environmental temperatures, but that effect reversed as temperatures increased. In contrast, skeletonizing herbivory declined with increasing functional and phylogenetic diversity, but only at low elevation sites, characterized by high environmental temperatures. Consequently, the net effect of elevation on herbivory varied by herbivore guild, with increasing elevation causing a net increase in chewing, a net decrease in skeletonizing, and no overall impact on herbivory caused by thrips, leaf mining, or galling (Fig 5).

**Figure 5.**
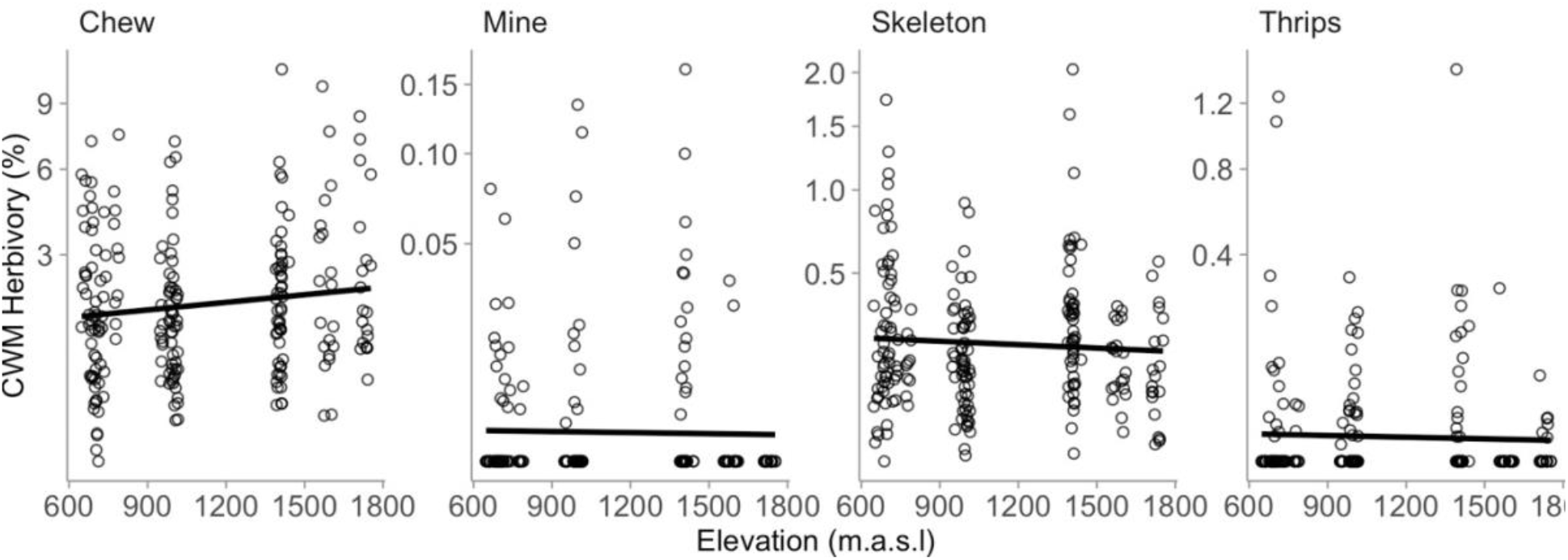
Community-weighted mean herbivory as a function of increasing elevation in the CBO divided into four herbivore functional groups.

Together, these results suggest that the effects of changing environmental temperature on invertebrate herbivory cannot be isolated from the biological context in which they take place. Specifically, in contrast with predictions grounded in theory and past experimental studies, our results show that the effects of changing environmental temperatures on herbivory depend on characteristics of plant communities, and how those plant communities modify abiotic drivers of herbivory, in turn, depend on feeding strategies of the herbivores under consideration. Consequently, these results highlight the need to consider characteristics of herbivores and the plant communities that they occupy when predicting how herbivory and herbivore communities will respond to increasing environmental temperatures associated with climate change.

## Discussion

This study shows, to our knowledge, the first field evidence that the abiotic environment can govern whether and how dimensions of plant biodiversity impact invertebrate herbivores. Not only can abiotic conditions directly influence invertebrate herbivory, the abiotic environment can also indirectly influence herbivory by simultaneously driving variation in plant diversity itself (mediation) (e.g., Warner *et al*. 2021), and by modifying how plant diversity impacts invertebrate herbivores (moderation) (e.g., Hülber *et al*. 2015; Field *et al*. 2020). Furthermore, these responses are strongly sensitive to invertebrate herbivore feeding guilds (e.g., Andrade *et al*. 2020), suggesting that any single predictor was inadequate for explaining herbivory along our environmental gradient. Together, these results reveal an important role that plant communities can play in driving herbivory across environmental gradients, suggesting that predicting how climate change will influence herbivory will require explicit consideration of how plant and herbivore communities respond to global change.

Counterintuitively, our results indicate that increasing temperature, associated with lower elevations can reduce invertebrate herbivory. When we explored this pattern in the context of herbivore feeding guilds, we found that this effect was restricted to chewing herbivores. This result is consistent with the hypothesis that increasing temperature should most strongly impact invertebrates with free-living life stages that feed externally on leaf tissue (Anstett *et al*. 2014), which has been invoked to explain why leaf chewing insects experienced a substantially stronger response to increasing annual rainfall compared to other invertebrate feeding guilds in a seasonally dry tropical forest (Andrade *et al*. 2020). These results further corroborate past studies suggesting that environmental gradients like temperature can directly alter the strength of biotic interactionslike herbivory, predation, or symbioses (Schemske *et al*. 2009; Rasmann *et al*. 2014; Descombes *et al*. 2017; Roslin *et al*. 2017; Hargreaves *et al*. 2019).

In addition to directly influencing herbivory, our results indicate that changing environmental temperatures can indirectly influence herbivory by altering multiple dimensions of plant community diversity. Specifically, increasing temperature (associated with lower elevations) reduced plant community phylogenetic diversity, which, in turn, increased herbivory (though only at higher temperatures). This shift in phylogenetic diversity might be attributable to the occurrence of both low-elevation and high-elevation adapted plant species at the highest study sites (Colwell & Lees 2000). Communities in the highest elevation meadow, located just below the tree line may represent an intermediate zone between subalpine and alpine vegetation communities, including plant species that are typically associated with lower and higher elevations (Halliday *et al*. 2021).

Our results further indicate that changing environmental conditions can modify the fundamental effects of host community diversity on invertebrate herbivory (moderation). Specifically, increasing plant functional diversity reduced herbivory, but only at low temperature, high elevation sites, and increasing plant phylogenetic diversity increased herbivory, but only at high temperature, low elevation sites. Changing temperature can alter plant defense compounds, palatability, and tissue nutrient concentrations, as well as the relative timing and rates of plant and herbivore growth and phenology (Andrade *et al*. 2020; Hamann *et al*. 2021; Kharouba & Yang 2021), so perhaps it is not surprising that functionally or phylogenetically diverse communities in one environment might not play the same role in determining herbivory in another environment. This dependence of community-level drivers of herbivory on environmental conditions has been observed in other systems as well. For example, in a study with a single focal plant species, increasing plant diversity reduced herbivory, and that effect was stronger in experimentally irrigated communities (Field *et al*. 2020). In a study of Swiss alpine grasslands, community-level herbivory increased with increasing species richness, soil fertility, and plant abundance, and that effect was stronger at higher elevation (characterized by cooler environmental temperatures), even though the total amount of herbivory was lower (Hülber *et al*. 2015). In an experimental grassland, invertebrate herbivory increased with increasing plant species richness (but not functional diversity), and this effect was stronger in summer than in spring (Meyer *et al*. 2017). Together, these results suggest that a temperature increase can shift the balance between different diversity effects on herbivory from negative (e.g. associational resistance) to positive (e.g. associational susceptibility, dietary mixing) (Bernays *et al*. 1994; Barbosa *et al*. 2009; Underwood *et al*. 2014) (but see Pellissier *et al*. 2012) and that these effects might depend on which dimensions of diversity are considered under which environmental conditions.

These statistical interactions between environmental temperature and dimensions of plant community diversity were themselves sensitive to differences among herbivore feeding guilds: chewing and skeletonizing herbivores had strong and opposing relationships between all three dimensions of diversity, temperature, and herbivory, while other feeding guilds had weak or undetectable responses. Specifically, increasing taxonomic, functional, and phylogenetic diversity tended to reduce chewing herbivory at low temperature, while that effect weakened (and reversed for functional and phylogenetic diversity) as temperature increased. In contrast, increasing taxonomic, functional, and phylogenetic diversity tended to reduce skeletonizing herbivory, but only at high temperature. A large body of literature suggests that responses to biotic and abiotic factors should differ among feeding guilds (Tscharntke & Greiler 1995; Bale *et al*. 2002; Anstett *et al*. 2014; Andrade *et al*. 2020), and yet to our knowledge, this is the first study exploring how these three factors (the biotic environment, abiotic environment, and herbivore feeding guilds) interact with one another.

Our results may explain why past studies have observed inconsistent responses of invertebrate herbivory to changing environmental temperatures in natural ecosystems (Moles *et al*. 2011; Galmán *et al*. 2018; Zvereva *et al*. 2020). Previous studies have observed dramatic reductions in herbivory with increasing elevation (e.g., Pellissier *et al*. 2014) or latitude (e.g., Baskett & Schemske 2018), or remarkably consistent increases in herbivory with increasing elevation (e.g., Sethi & Hille Ris Lambers 2021) and latitude (e.g., Adams & Zhang 2009). In experiments, herbivore richness, abundance, community structure, and overall rates of herbivory are all strongly influenced by plant species richness (Haddad *et al*. 2001; Scherber *et al*. 2006; Ebeling *et al*. 2018). Other characteristics of the plant community are important in shaping the composition of arthropod communities as well (Gardarin *et al*. 2018), which can feed back and drive community level herbivory (Fernandez-Conradi *et al*. 2022). Our results suggest that some of the inconsistent patterns observed in previous observational studies might stem from non-independence among these biotic and abiotic drivers of herbivory. However, we cannot rule out the possibility that the effects observed along this elevation gradient were the result of unmeasured processes associated with elevation, including unmeasured abiotic variables, feedbacks between top-down and bottom-up regulation of herbivores, and adaptation of plants and herbivores to their local biotic and abiotic environment. Future studies could address these research gaps using combinations of reciprocal transplant, common garden, and warming experiments along elevational gradients (Buckley *et al*. 2019; Zvereva *et al*. 2022).

Together, the results of this study highlight a growing need to consider plant and herbivore community context in order to predict how climate change will alter invertebrate herbivory. Specifically, in this study, the most important drivers of herbivory were multiple dimensions of plant diversity, but how those dimensions of diversity impacted herbivory depended on both the abiotic conditions and on the feeding guilds of herbivores under consideration. These results support a growing body of literature suggesting that how plant communities regulate ecosystem processes may depend on characteristics of species present in those ecosystems (Lavorel & Garnier 2002; Mouillot *et al*. 2011; Allan *et al*. 2015; Leitão *et al*. 2016; Funk *et al*. 2017; Van de Peer *et al*. 2018; Le Bagousse-Pinguet *et al*. 2019; Heilpern *et al*. 2020), but that abiotic factors such as temperature can override the effects of biotic factors on ecosystem processes (Cannone *et al*. 2007; Laiolo *et al*. 2018; Halliday *et al*. 2021). These results therefore suggest that predicting how global change will influence herbivory may depend on complex relationships between global change drivers and the structure of plant and herbivore communities.

## Materials and methods

### Study system

The Calanda Biodiversity Observatory (CBO) consists of five publicly owned meadows located along a 1101 m elevational gradient (648 m to 1749 m) below tree-line on the south-eastern slope of Mount Calanda (46°53′59.5″N 9°28′02.5″E) in the canton of Graubünden (Halliday *et al*. 2021). The soil in the area is generally calcareous and has low water retention (Eggenberg & Möhl 2013; Alexander *et al*. 2015). The five CBO meadows are variable in size (roughly 8 to 40 Ha), and separated by forests and at least 500 m elevation. Meadows are maintained through grazing and mowing, a typical form of land use in the Swiss Alps (Bätzing 2015), and cover collinean (<800 m) mountain (800 m–1500 m) and subalpine (1500–2200 m) vegetation zones (Ozenda 1985; Eggenberg & Möhl 2013). The CBO meadows are grazed by cattle twice per year as the cattle are moved between low and high altitudes.

The CBO consists of a nested set of observational units. Each meadow contains 2–7, .25 ha sites (n = 22 sites), each of which contains a grid of nine evenly spaced, 4m^2^ large-plots, with the exception of one site (I3), which is 100 m x 25 m and contains 10 large plots due to its shape (n = 199 large plots). In each site, large plots are arranged in a grid with the center of each plot separated by at least 20m distance from its nearest neighbor. Each large plot is subdivided into four, 1 m^2^ subplots (n = 796). At each site, five large plots were selected to contain an intensively surveyed module, which consisted of two 50 cm-diameter, round small plots, placed in opposite subplots (n=110 intensively surveyed modules consisting of 220 small plots). These intensively surveyed small plots are the smallest unit of observation used in this study.

### Quantification of plant community diversity

In July 2019, we recorded the identity and visually quantified the percent cover of all plant taxa in each small plot following a modified Daubenmire method (n = 220) (Halliday *et al*. 2021). Using these data, we evaluated changes in three dimensions of plant community diversity to evaluate indirect effects of environmental conditions on herbivory: plant species richness, plant functional divergence, and plant phylogenetic overdispersion. These three components of host community structure commonly respond to changing environmental conditions (Hulshof *et al*. 2013; Descombes *et al*. 2017), and represent important characteristics of plant communities that can influence herbivory. We quantified functional divergence (FDiv; Villéger *et al*. 2008), using the dbFD function in R package FD (Laliberté *et al*. 2014), as a measure of functional diversity that is independent of taxonomic and phylogenetic diversity. Using the TRY database (Kattge *et al*. 2020), we extracted nine traits for every plant taxon (specific leaf area, carbon-to-nitrogen ratio, leaf chlorophyll content, leaf lifespan, leaf nitrogen, leaf phosphorus, and photosynthetic rate, plant height, and seed mass), omitting tree seedlings, which are functionally dissimilar from the more dominant herbaceous taxa, and taxa that could not be identified to host genus, which together, never accounted for more than 7% cover in a plot (mean = 0.04%). Unknown taxa that could be identified to the genus level were assigned genus-level estimates for each trait, by taking the mean of the trait value for all members of that genus that had been observed on Mount Calanda during extensive vegetation surveys.

We computed phylogenetic overdispersion to quantify phylogenetic diversity independent of species richness and functional diversity, by constructing a phylogeny of all non-tree species using the phylo.maker function in R package V.PhyloMaker (Jin & Qian 2019). Plant phylogenetic diversity for each small plot was calculated using the ses.mpd function in R package Picante (Kembel *et al*. 2010). This function uses a null-modeling approach that measures the degree to which a small plot is more or less phylogenetically diverse than random, given the number of host species, weighted by their relative abundance. Specifically, we generated a z-score that compares the mean-pairwise-phylogenetic-distance between taxa in a plot to a randomly assembled plot with the same number and relative abundance of host species, drawn from the full species pool, and permuted 1000 times.

### Quantification of herbivory

A survey of foliar herbivory was carried out in August 2019 by estimating the percent of leaf area damaged by invertebrate herbivores on up to five leaves of twenty randomly selected plants per small plot (n = 18203 leaves on 4400 plants across 220 small plots). This survey was carried out by placing a grid of 20 equally spaced grill sticks into the ground, with each stick having a distance of 10 cm to its nearest neighbor. The 20 plant individuals that were most touching the sticks were then identified, and the five oldest non-senescing leaves on each plant were visually surveyed for herbivory following the plant pathogen and invertebrate herbivory protocol in Halbritter et al (2020). Some plant individuals had fewer than five leaves, so fewer than five leaves were surveyed on those plants.

Herbivory was assessed for each small plot using the community weighted mean leaf area damaged by herbivores, calculated as the mean leaf area damaged by herbivores on a plant species in a plot, multiplied by the relative abundance of that plant species from the vegetation survey, and then summed across all plant species in the plot.

### Quantification of environmental conditions

As described in Halliday et al (2021) soil temperature (6 cm below the soil surface), soil surface temperature, air temperature (12 cm above the soil surface), and soil volumetric moisture content were recorded at 15 minute intervals for 22–37 days (average 31 days) in the central large plot of each site (n = 22) using a TOMST-4 datalogger (Wild *et al*. 2019).

### Statistical analysis

All statistical analyses were performed in R version 3.5.2 (R Core Team 2015). All analyses consisted of fitting linear mixed models with an identity link and Gaussian likelihoods using the lme function in the nlme package (Pinheiro *et al*. 2016). In order to meet assumptions of normality and homoscedasticity, we square-root transformed community weighted mean herbivory and added an identity variance structure (varIdent function) for each site, which based on visual inspection of residuals of each model, exhibited considerable heteroscedasticity (Zuur *et al*. 2009; Pinheiro *et al*. 2016). Each model included large plots, sites, and meadows as nested random intercepts to account for non-independence among observations due to the sampling design of the CBO. Full equations and parameters for these models are available on Github (https://github.com/fhalliday/Calanda19_Herbivory).

To compare direct and indirect effects of environmental conditions on herbivory, we performed confirmatory path analysis using the PiecewiseSEM package (Lefcheck 2016). Specifically, we fit a structural equation model (SEM) that included the effect of elevation on air temperature, the effect of air temperature on square-root-transformed herbivory, the effect of air temperature on three endogenous mediators (plant taxonomic, functional, and phylogenetic diversity), and the effects of those two mediators on square-root-transformed herbivory. We also tested the hypothesis that air temperature could alter the relationship between plant community diversity and herbivory by fitting a second-stage moderated mediation (Hayes 2015) including the pairwise interaction between air temperature and each endogenous mediator. To aid the interpretation of direct effects in the model, we mean-centered air temperature and each endogenous variable, so that average values were used as the reference values for interpreting the other variables’ independent effects. We then explored the interaction between each measure of plant community diversity and temperature by setting the reference temperature to one standard deviation above and below the mean temperature, and re-running the model.

To explore differences in the responses to biotic and abiotic drivers among herbivore feeding guilds, we fit a multiresponse regression model (following Dooley *et al*. 2015; Halliday *et al*. 2017). This model included four dependent variables: square-root transformed community-weighted-mean herbivory caused by chewing, leaf mining, skeletonizing, and thrips. We standardized the response variables to the same scale by dividing each observation by the maximum value for that response. To model whether the effects differed among these responses, we constructed our multiresponse regression with mean air temperature, plant taxonomic diversity, plant functional diversity, and plant phylogenetic diversity as fixed effects. To estimate whether the effect of plant community structure depend on temperature, we also included in the model the pairwise interactions between each of the three measures of host community structure and temperature as additional fixed effects.

## Data accessibility statement

The data and code supporting the results are available on Github (https://github.com/fhalliday/Calanda19_Herbivory).

## Acknowledgements

We are grateful for insightful suggestions and field assistance from M. Jalo, J. Alexander, M. Maechler, K. Raveala, M. Tiusanen, A. Norberg, I. Kohonen, V. Loaiza, J. Moser, and members of the Laine Lab. This work was supported by Gemeinde Haldenstein, Gemeinde Chur, the University of Zürich, and by grants from the Academy of Finland (296686) to A-LL, the European Research Council (Consolidator Grant RESISTANCE 724508) to A-LL, and an Ambizione Grant (PZ00P3_202027) from the Swiss National Science Foundation to FWH.

---

**Figure S1.**
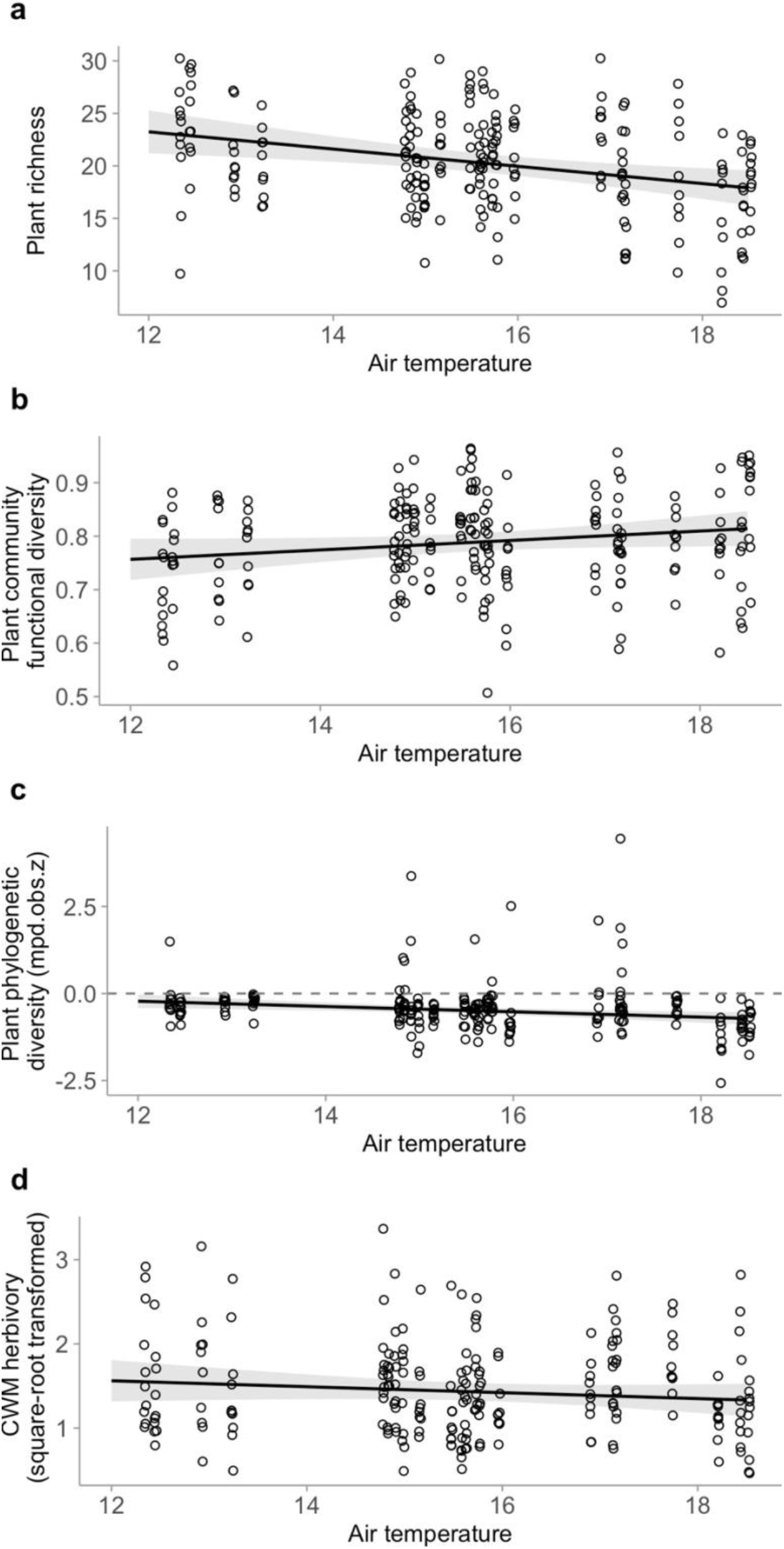
Results of linear models predicting (a) taxonomic, (b) phylogenetic, (c) functional diversity, and (d) community-weighted mean (CWM) herbivory as a function of increasing air temperature.

**Figure S2.**
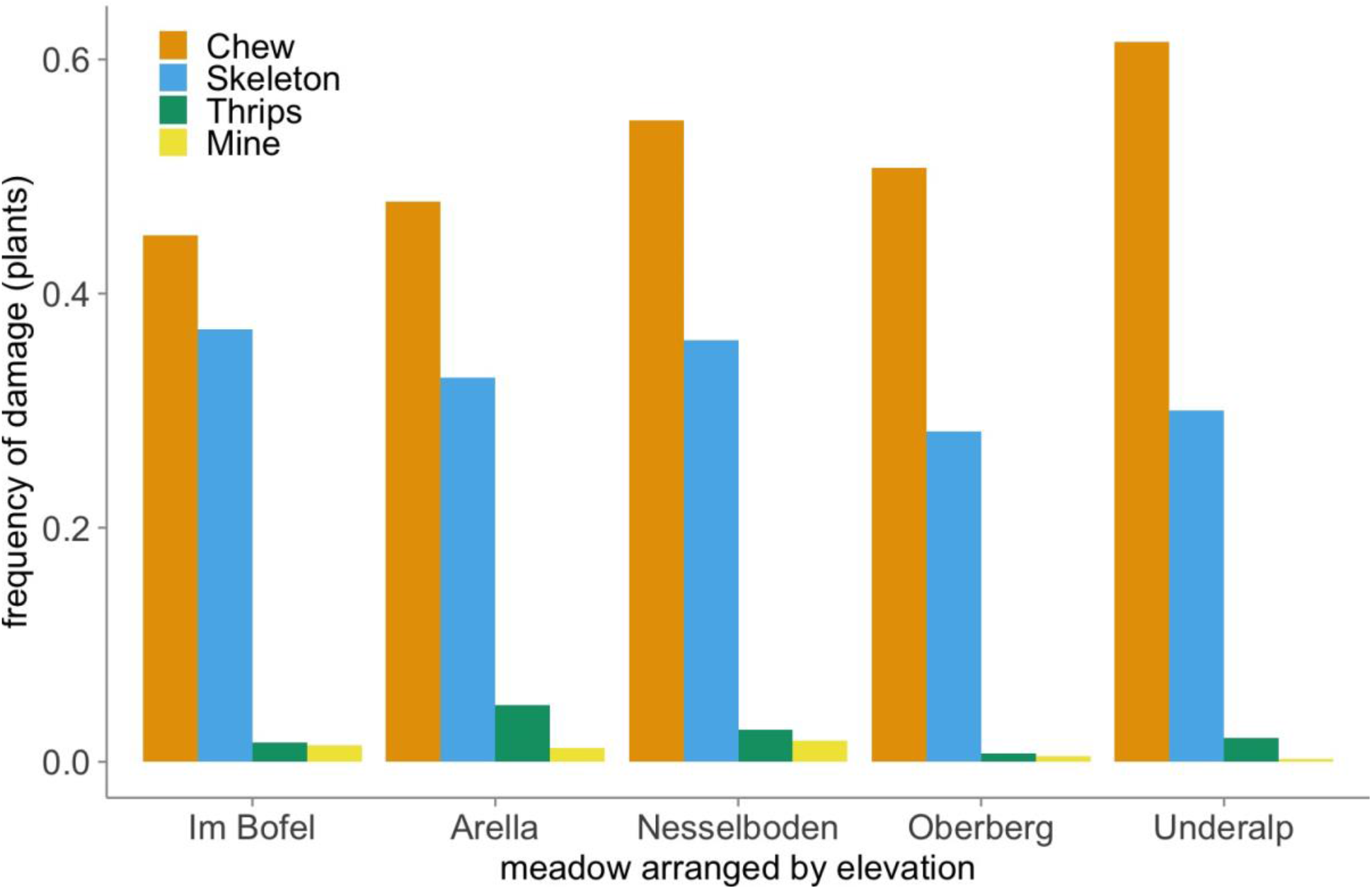
The percent of plants damaged by each feeding guild in each meadow. Meadows are arranged from lowest elevation (~600 m.a.s.l) to highest elevation (~1700 m.a.s.l).

**Table S1.** Results of Type II Analysis of Deviance tests on models quantifying the effects of air temperature on three measures of host community structure.

| Response | Chisq | Df | P |
| --- | --- | --- | --- |
| Host Richness | 10.36 | 1 | 0.0012 |
| Host Functional Diversity | 3.185 | 1 | 0.0743 |
| Host Phylogenetic Diversity | 8.814 | 1 | 0.0030 |

**Table S2.** Coefficient estimates from the structural equation model. Estimates are provided both raw and standardized to a common scale to facilitate comparisons

| Response | Predictor | Estimate | Std Error | DF | Critical Value | P | Std Estimate |
| --- | --- | --- | --- | --- | --- | --- | --- |
| Square-root transformed community-weighted mean herbivory | Air temperature | -0.4469 | 0.1880 | 16 | -2.3771 | 0.0303 | -1.4134 |
|  | Species richness | -0.0138 | 0.0076 | 104 | -1.8223 | 0.0713 | -0.1138 |
|  | Functional diversity | -0.7250 | 0.3894 | 104 | -1.8620 | 0.0654 | -0.1118 |
|  | Phylogenetic diversity | 0.0044 | 0.0542 | 104 | 0.0816 | 0.9351 | 0.0057 |
|  | Temperature × Richness | -0.0064 | 0.0042 | 104 | 1.5233 | 0.1307 | 0.4174 |
|  | Temperature × Functional | 0.4078 | 0.2031 | 104 | 2.0082 | 0.0472 | 1.0072 |
|  | Temperature × Phylogenetic | 0.0764 | 0.0353 | 104 | 2.1669 | 0.0325 | 0.1964 |
| Species richness | Air temperature | -0.8213 | 0.2552 | 16 | -3.2186 | 0.0054 | -0.3155 |
| Functional diversity | Air temperature | 0.0088 | 0.0049 | 16 | 1.7846 | 0.0933 | 0.1812 |
| Phylogenetic diversity | Air temperature | -0.0763 | 0.0257 | 16 | -2.9687 | 0.0091 | -0.1861 |
| Air temperature | Elevation | -0.0051 | 0.0005 | 214 | -9.5685 | 0.0000 | -1.0430 |

Goodness of fit: Fisher's C = 13.143 with p = 0.515 and on 14 degrees of freedom

